# Automated Image-Based Cell Sorting by Targeted Photopolymerization

**DOI:** 10.64898/2025.12.27.693999

**Authors:** Stefan J. Maurer, Tobias Abele, Saskia R. Stange, Kira H. Hoffmann, Isabel Poschke, Michael Platten, Kerstin Göpfrich

**Affiliations:** Biophysical Engineering Group, Center for Molecular Biology of Heidelberg University (ZMBH), Berliner Str. 45, Heidelberg, D-69120, Germany; Clinical Cooperation Unit (CCU) Neuroimmunology and Brain Tumor Immunology, German Cancer Research Center (DKFZ), Im Neuenheimer Feld 280, Heidelberg, D-69120, Germany; German Cancer Consortium (DKTK), DKFZ, core center, Heidelberg, Germany; Immune Monitoring Unit, National Center for Tumor Diseases (NCT), NCT Heidelberg, a partnership between DKFZ and Heidelberg University Hospital, Heidelberg, Germany; Faculty of Biosciences Heidelberg University Heidelberg, Germany; Department of Neurology, Medical Faculty Mannheim, Mannheim Center for Translation Neuroscience (MCTN) Heidelberg University, Mannheim, Germany; Helmholtz Institute for Translational Oncology Mainz (HI-TRON Mainz) – A Helmholtz Institute of the DKFZ, Mainz, Germany; DKFZ Hector Cancer Institute at the University Medical Center Mannheim, Mannheim, Germany

## Abstract

We present an automated image-based cell sorting method capable of a throughput of hundreds of cells per second. Our microscopy platform integrates automated image acquisition, machine-learning based classification and subsequent depletion of up to 99.98 % of negative cells by spatially controlled photo-encapsulation, preserving target cells for collection. Applied to peripheral blood mononuclear cells, the method achieves the label-free, morphology-based enrichment of activated T cells for downstream applications in biomedicine.

---

Cell sorting is required for fundamental research, analysis and therapy with applications ranging from functional and phenotypic characterization to enrichment for adoptive transfer for cell therapy. The most widely known method is fluorescence activated cell sorting (FACS) [1] but image-based methods gain traction as their spatio-temporal resolution permits label-free sorting according to cellular morphology and sub-cellular features [2].

Image-based sorting methods can be divided into two sub-categories. For one, microfluidic flow platforms, in which cells traverse a channel in which they are probed and a sorting decision is carried out downstream [3–8], and stationary imaging platforms, where cells are plated on address sites and often immobilized for imaging [9–15]. Stationary methods provide high quality sorts thanks to specialized imaging without the need for complex corrections due to motion blur [16] and offer the possibility of observing individual cells over extended periods of time [2]. However, these advantages come at the cost of low throughput and often the need for specialized equipment, expert knowledge, tedious sample preparation, and high equipment cost.

Here, we introduce an automated and versatile microscope-based platform for image-based sorting that carries the advantages of platform-based approaches but with improved throughput. At the core of our technique lies the selective encapsulation of cells in a photopolymerizable solution (photoresist) supplemented to the cell medium. Encapsulated cells can be subsequently filtered out of the suspension, thereby isolating non-encapsulated cells that carry the desired feature. Our setup fully integrates hardware components, namely a microscope and a light-emitting diode (LED) illuminated digital micromirror device (DMD), and our custom software that guides the user through the automated sorting process.

In particular, cells are sorted in four steps as shown in Figure 1. Cells are first resuspended in cell medium containing a cytocompatible photoresist, which polymerizes upon millisecond illumination with an LED at 385 nm wavelength. For sorting, the suspension is then inserted into a customized 35 mm Petri dish-based chamber, in which the cells settle homogeneously. The chamber is placed under a microscope equipped with a DMD, which is integrated into its light path through an external camera port. The DMD projects light patterns onto the sample to initiate localized photopolymerization of the photoresist and thus selective encapsulation of the targeted cells. We developed a Python-based software package that controls the microscope and DMD, performs image analysis, and guides the user through the encapsulation process *via* an intuitive graphical user interface. Here, the user can select the hardware configuration for imaging the sample, such as the objective or the imaging mode (bright-field / fluorescence) as for any conventional microscopy imaging experiment, and set up the image analysis. To this end, the software provides a modular framework that assists the user in designing an analysis pipeline, leading to the classification of cells imaged by the microscope. Predefined analysis pipelines are available, but custom Python-based analysis functions can be easily integrated into the framework to adapt the method to the application at hand.

**Fig. 1:**
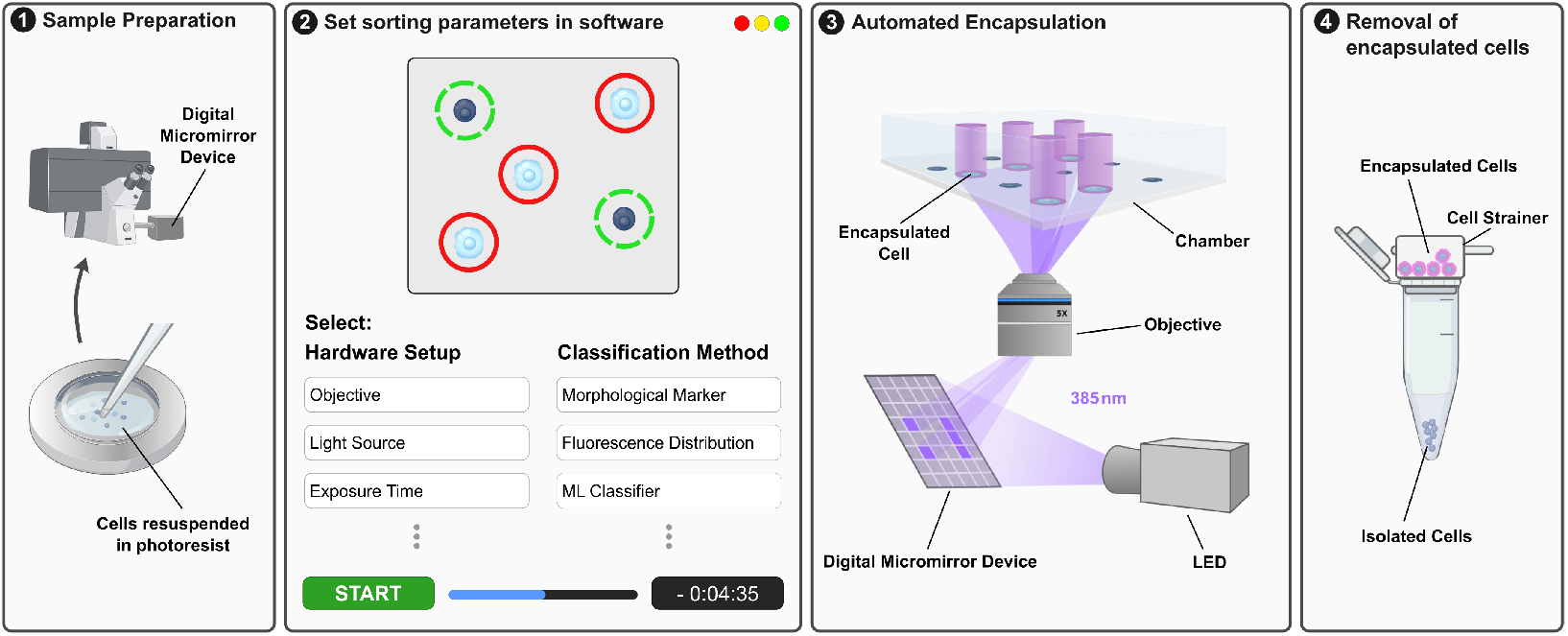
Workflow for image-based cell sorting by automated, targeted photopoly-merization. **1** Cytocompatible photoresist is added to the culture media and the cell suspension filled into a chamber, which is placed under a microscope equipped with an LED illuminated digital micromirror device (DMD). **2** The sorting parameters are configured in our custom software. This includes the microscope setup to image the cells and configuration of the analysis pipeline to detect and classify cells. **3** The user initiates the automated encapsulation process: The software first images and classifies cells tile by tile, then calculates photomasks, which are projected onto each tile by the DMD. This causes the photoresist in the cell medium to polymerize around the targeted population of cells only, thereby selectively encapsulating them. **4** The suspension is removed from the chamber and filtered through a cell strainer, which holds back encapsulated cells, isolating the cells of interest in the filtrate.

After the software has scanned and analyzed the sample, it computes binary masks from the positions of cells that need to be encapsulated. These masks are then projected by the DMD onto the sample to locally initiate photopolymerization around the cells across the entire chamber. To compensate for intensity falloff toward the edge of the projected area and ensure homogeneous photopolymerization across the entire field of view, we compute a correction map for the exposure time to adjust it individually for each micromirror of the DMD. After cell encapsulation, the entire solution is removed from the chamber and passed through a cell strainer; the encapsulated negative cells are retained and the filtrate containing the unencapsulated positive cells is collected.

Rate of recovery, depletion and throughput of the sorting method were bench-marked using fixed Madin-Darby Canine Kidney (MDCK) cells, which were stained separately with amine-reactive dyes, either fluorescein (green) or rhodamine X (orange). The cells were mixed in a 1:1 ratio and the mixture of green and orange cells was resuspended at different concentrations in a fast polymerizing photoresist based on poly (ethylene glycol) diacrylate (PEGDA) monomers, which we activated using a light-emitting diode at 385 nm wavelength. The cells were then classified according to their fluorescence signal and cells of one stain, here fluorescein, selectively encapsulated (Fig. 2a). By adjusting the software sorting parameters, as well as the cell density and repeat cycles, the setup can be optimized to prioritize the recovery rate, throughput, or depletion, respectively (Fig. 2b). A higher cell density reduces the spacing between cells and increases the number of cells that can be classified per frame, thereby increasing throughput. At the same time, a higher cell density increases the probability that positive and negative cells are too close in proximity to avoid co-encapsulation. Proximity-induced co-encapsulation of positive with negative cells reduces the recovery rate (Fig. 2c), counteracting the gain in throughput from increased cell density (Fig. 2d). The largest number of cells was reliably recovered at a cell density of ~200 000 cells per chamber with a recovery rate of 47 % when screening at a throughput of 500 cells/s. The rate of depletion of unwanted cells decreased slightly with higher cell density with an average of (99.1 ± 0.4) % (mean ± s.d., *n* = 17) (Fig. 2e).

**Fig. 2:**
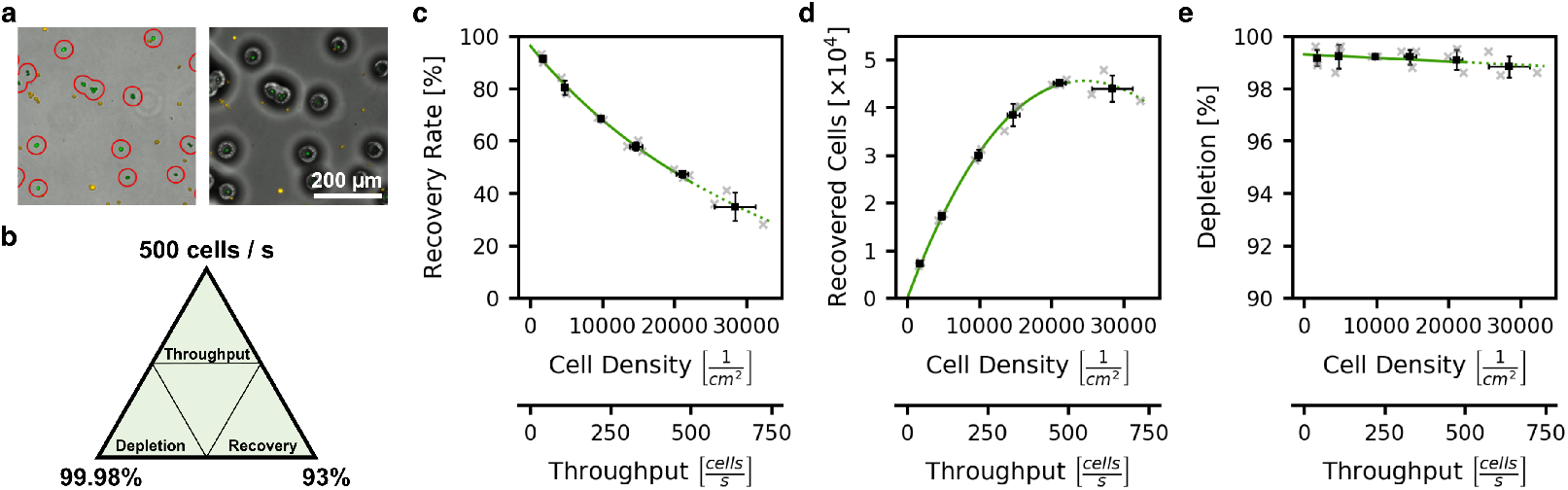
Benchmarks of DMD-assisted image-based sorting by photopolymerization. **a** For benchmarking, fixed MDCK cells were separately stained with either fluorescein (green) or rhodamine X (orange), mixed, and sorted according to their staining. Micrographs show cells before (left) and after (right) photo-induced encapsulation. **b** Maximum performance when the set up was optimized for either throughput, recovery or depletion, respectively. **c-e** Detailed benchmarking of recovery rate, recovered cells and depletion as functions of the cell density in the chamber, which is proportional to the throughput. **c** The recovery rate of positive cells decreased with increasing cell density, as the reduced intercellular distance between positive and negative populations increased the probability of co-encapsulation. **d** Number of recovered positive cells from a single sorting run. A maximum number of cells could be reliably recovered at a cell density corresponding to 500 cells/s. For exceeding cell densities (dotted line) large sections of the photoresist had to be polymerized impeding overall cell recovery. **e** Depletion showed a slight, approximately linear decrease with increasing throughput, averaging (99.1 ± 0.4) % (mean ± s.d., *n* =17). Function fit (green): Recoveryrate 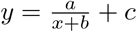, recovered cells 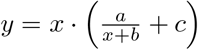, linear depletion.

In general, imaging at higher magnification, i.e. with a smaller field of view, reduces throughput. For example, if imaging magnification is increased from 5× to 10×, 20×, or 40×, the sorting time should increase by a factor of four, sixteen, or sixty-four respectively. While an entire chamber filled with cells can be sorted within 5 min at 5×, sorting at 20× would thus take 80 min. To compensate for the speed loss, we developed a scanning pattern that divides the sample into sections. Each section is first imaged in its entirety using a higher magnification objective, before negative cells within that section are encapsulated using the objective with the largest field of view available. At 5×, we were able to achieve a sufficiently high resolution to photopolymerize encapsulating structures with a diameter of ~ 50 µm with the DMD. When a higher magnification objective is selected for imaging and lower magnification objective is used for photopolymerization, our software is able to adjust the patterns of illumination accordingly. With this strategy, imaging at 5×, 10× and 20× resulted in a sorting time of 5 min, 9.5 min and 25 min per chamber, respectively. The ability of our method to encapsulate hundreds of cells in parallel can significantly speed up (by over two orders of magnitude for a 1:1 mixture of positive and negative cells) the sorting step compared to methods that target and retrieve cells individually, for example with a micropipette [10].

In the field of immunology, methods to isolate immune cell subsets are often required to understand cell-type-specific functions, behaviors, and molecular profiles, enabling insights into disease mechanisms and therapeutic targets [17]. In addition, such methodologies are often required for enrichment for immune cells, such as T cells for therapeutic purpose. For instance, the *ex vivo* genetic engineering of chimeric antigen receptor (CAR) T cells requires the isolation of large amounts of functional patient-derived T cells from peripheral blood [18]. This isolation is crude based on cell lineage-defining cell surface receptors. The resulting cell population is highly heterogeneous with respect to functionality and differentiation state. T cells are known to undergo morphological changes upon activation, for example an increase in cell volume [19]. To demonstrate the label-free, image-based capabilities of our system, we enriched *in vitro*-expanded, activated T cells from a mix with peripheral blood mononuclear cells (PBMCs). To this end, *in vitro*-expanded, activated T cells [20] and unstimulated PBMCs, which could be morphologically distinguished by their size, were mixed in different ratios and resuspended in the DexGMA-based photoresist (Fig. 3a). We developed a fast detection method for our software to identify cell contours from bright-field images and detected cells were classified by the equivalent circular diameter calculated from the contour area. Cells above a threshold of 11.5 µm were considered activated and thus positive cells, while cells below the threshold were encapsulated (Fig. 3b,c). Structures smaller than 5 µm were considered cell debris. To verify the selection by size, we added human embryonic kidney 293 (HEK 293) cells with a cell diameter of 11 µm to 15 µm to the mixtures, which confirmed enrichment (Fig. 3d). Additionally, sorting resulted in an increased proportion of myeloid cells from the unstimulated PBMCs, as e.g. larger monocytes have diameters exceeding 11.5 µm [21]. We validated the enrichment of activated T cells post sorting by comparing the level of T cell activation marker CD69 in CD3+ cells in the pre- and post-sorting samples (Fig. 3e). The cell viability remained high, declining only marginally, throughout the sorting process (98.7 % to 99.5 % pre and 92 % to 98.7 % post sorting).

**Fig. 3:**
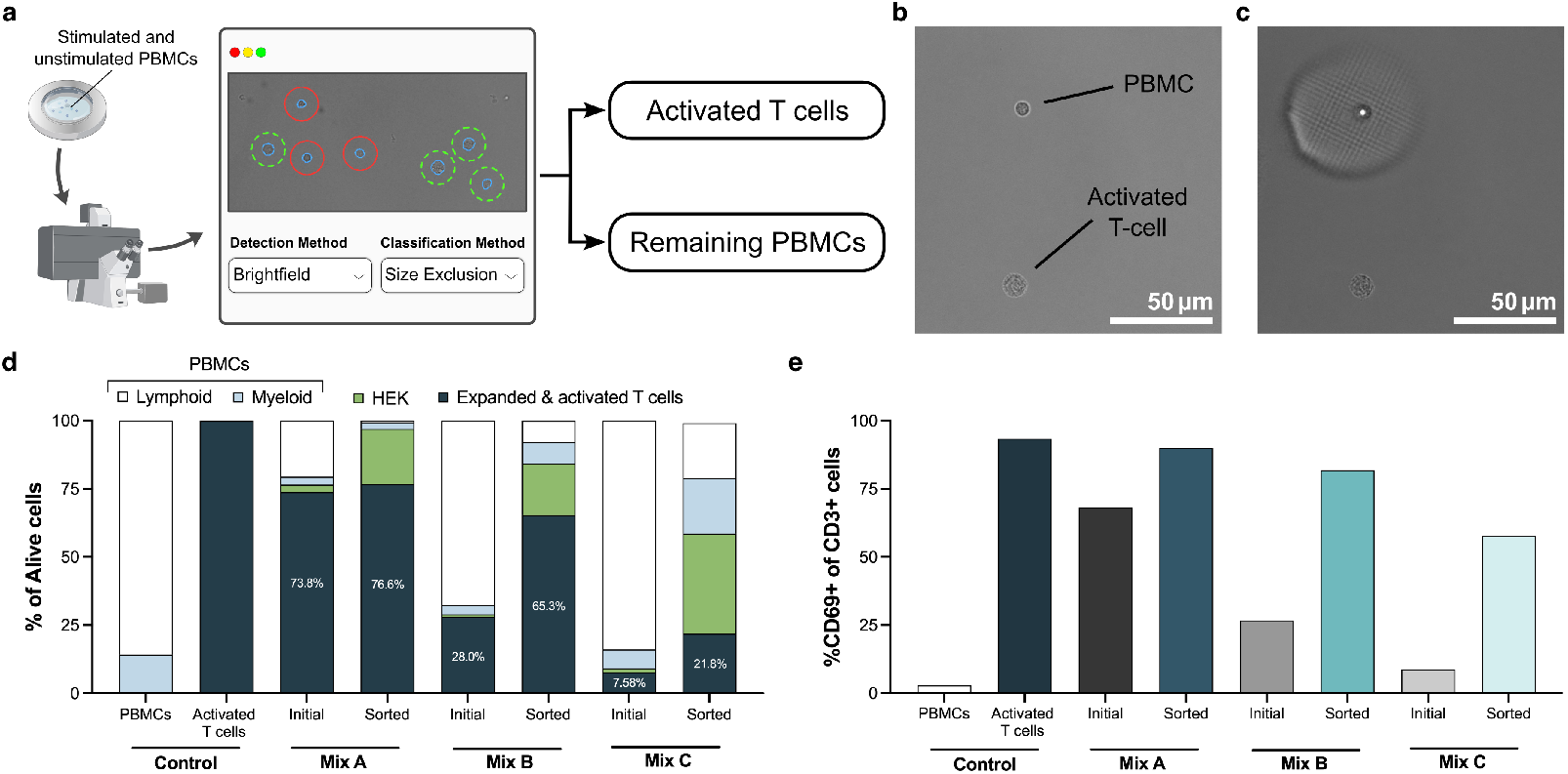
Label-free, image-based isolation of *in vitro*-expanded, activated T cells from a mixture with unstimulated PBMCs. **a** Contours (blue) of cells and cell size are detected from bright-field images through modified edge-detection. To isolate the *in vitro*-expanded, activated T cells, cells with a diameter of 5 µm to 11.5 µm are selectively encapsulated, and subsequently removed, retaining large activated cells. **b** *In vitro*-expanded, activated T cell and unstimulated PBMC before and **c** after encapsulation of cells with a diameter of 5 µm to 11.5 µm. **d** Validation of cell type-specific enrichment based on equivalent cell diameter by staining of *in vitro*-expanded, activated T cells (dark teal), HEK 293 cells (green) and unstained PBMCs separated as lymphoid (white) and myeloid (light blue) prior to mixing and sorting. **e** Comparing levels of T cell activation marker CD69 in CD3+ cells before and after sorting shows clear enrichment of activated T cells across mixtures with different initial concentrations.

We provide an easy-to-use platform that combines image- and fluorescence-based sorting from high quality images, with a higher throughput than comparable methods. Our method sorts cells directly in a custom Petri dish, eliminating the need for compartments or microarray address sites. By separating the classification and active sorting step, computationally expensive image analysis is possible in-between, without limiting throughput. We demonstrate the utility of our approach by sorting for T cell activation in a label-free manner based on bright-field imaging. Our platform can in principle be implemented on specialized microscopes, such as confocal or super-resolution microscopes, as the only requirement is a free external camera port for the attachment of a DMD-based pattern projector, thus enabling the selection of cells based on advanced imaging techniques. Importantly, our technology is compatible with sorting based on timelapse images, allowing users to observe phenotypical changes over time before making a sorting decision. In the future, the method could be extended to sort adherent cells, organoids, and cells in 3D culture. Furthermore, our approach has the potential to isolate synthetic cells and vesicles for applications in synthetic biology, biotechnology, and medicine.

## Materials and Methods

### Hardware setup

Sorting was carried out on an inverted widefield microscope (Axio Observer 7, Carl Zeiss AG) with a digital micromirror device (DMD) pattern illuminator (Polygon 1000, Mightex) attached through an external camera port. A light emitting diode (Mightex) emitting light with a wavelength of 385 nm was connected to the DMD through a light guide. The microscope and DMD were connected to a personal computer and controlled using custom software written in Python 3.11.12. Microscope control was made possible through integration of dynamic libraries from the Micro Toolbox Software Development Kit 3.2.11.0 provided by Zeiss into our Python application. Binary arrays, in which individual mirrors in the ‘on’ position are represented as ‘1’ and mirrors in the ‘off’ position are represented as ‘0’, could be transmitted to the DMD through a TCP/IP interface in the software provided by the manufacturer (Polyscan 4, Mightex). To compensate for intensity falloff toward the edge of the projected area and ensure even photopolymerization across the field of view, a per-micromirror exposure time correction map is applied on the DMD.

### Benchmark figure calculations

We need to consider sample loss when calculating the depletion *d* of the sample. Therefore, we calculate the depletion from the drop in odds of negative cells in the sample relative to their starting odds:

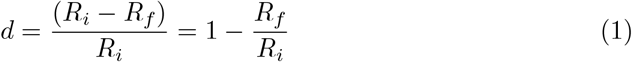

where *R*_*i*_ and *R*_*f*_ are the ratio of negative cells to positive cells before and after sorting, respectively.

The recovery rate *r* was calculated as:

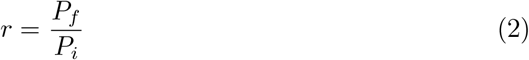

where *P*_*i*_ and *P*_*f*_ are the number of positive cells before and after sorting, respectively.

The throughput *t* was defined as:

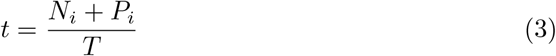

where *N*_*i*_ + *P*_*i*_ is the number of processed cells, and *T* is the time needed for imaging and encapsulation.

### Cell culture

After fixation in ice-cold ethanol under vortexing, MDCK cells were resuspended in Dulbecco’s phosphate-buffered saline (Gibco DPBS, ThermoFisher) with 2 wt% bovine serum albumin (BSA, Albumin fraction V, Carl Roth) for at least 12 h. The cells were washed twice in DPBS and stained with ROX NHS-Ester (Rhodamine X, Lumiprobe) or NHS-Fluorescein (5/6-carboxyfluorescein succinimidyl ester, Thermo Fisher Scientific) and stored in the fridge.

PBMCs and *in vitro*-expanded T cells of healthy blood donors were thawed, washed and rested overnight at 37 °C, 5 % CO2 in X-Vivo15 media (Lonza) containing 2 vol% human AB serum (Sigma-Aldrich). On the next day, cells were counted and seeded into a 6-well plate with 5 × 10^6^ cells/well in fresh media. For stimulation, *in vitro*-expanded T cells were cultured in media containing 50 µL/well TransAct anti-CD3/CD28 beads (Miltenyi). After 2 days, cells were harvested and counted. *In vitro*-expanded, activated T cells and HEK 293 cells were stained with 50 nM CellTrace Violet and 50 nM Cell-Trace CFSE (Life Technologies) diluted in Dulbecco’s phosphate-buffered saline (PBS, Sigma-Aldrich), respectively, according to manufacturer’s instructions. After staining, cells were resuspended in fresh Dulbecco’s Modified Eagle Medium (DMEM, Sigma-Aldrich) supplemented with 100 IU mL^*−*1^ penicillin and 100 µg mL^*−*1^ streptomycin (Life Technologies) and 10 vol% fetal bovine serum (FBS Superior, Sigma-Aldrich) to a final concentration of 1 × 10^6^ cells/mL and combined into different cell mixes.

### Sorting of fixed Madin-Darby Canine Kidney (MDCK) cells

For sorting, fixed MDCK cells in suspension stained with either Rhodamine X or Fluorescein were mixed in a 1:1 ratio, ensuring that the number of cells from each suspension was equal. The cells were resuspended in 450 µL Phosphate-buffered saline (PBS) with 2 wt% BSA, 30 vol% poly (ethylene glycol) diacrylate (PEGDA) and 15 g L^*−*1^ lithium phenyl-2,4,6-trimethylbenzoylphosphinate (LAP). A sorting chamber was built out of the parts of a 35 mm tissue culture-treated polystyrene petri dish (Falcon). The lid was inverted and repurposed as the bottom of the sorting chamber. A hole with a diameter of 2 mm was drilled into the center of the petri dish itself, which was then stacked onto the inverted lid, creating a narrow chamber in-between. The cell suspension pipetted into the chamber through the hole in the center would spread in the narrow spacing between bottom and lid and the cells would settle homogeneously spaced out on the entire surface of the bottom part.

The chamber was placed under the microscope and cells allowed to settle for 20 min. The entire sample was imaged in the ZEN Software (Zeiss) using the ZEN Macro Environment to count the cells. Our custom software then imaged the sample tile by tile either detecting ROX or Fluorescein stained cells using the 5× objective (Plan-Apochromat 5x/0,16 M27, Carl Zeiss AG) and identified cell positions from the images by thresholding and contour detection. After imaging each tile, the appropriate light-pattern was calculated by the software from all cell positions detected in the chosen channel and these cells were encapsulated immediately with an exposure time of 300 ms.

After encapsulation, the lid was removed from the chamber, the cells rinsed out using PBS with 2 wt% BSA and polymerized structures filtered using a 40 µm mesh cell strainer (Greiner Bio-One). The filtrate was collected in a 15 mL Falcon tube (Sarstedt AG & Co. KG) and centrifuged at 1000 rpm for 10 min before the supernatant was removed except for the bottom 1 mL of suspension, which was imaged in a 6-well-plate (Techno Plastic Products AG). For the high depletion sorting runs, the release of three chambers was sorted again in the same way.

Cells were counted with a FIJI [22] macro, using thresholding and the Analyze Particles plug-in.

### Sorting of peripheral blood mononuclear cells (PBMCs)

A custom glass-bottom chamber was built in the following way: A hole 2 mm in diameter was drilled into the center of a 35 mm tissue culture-treated polystyrene petri dish (Falcon) and a ~ 1 cm high ring was 3D-printed (X1-Carbon, Bambu Lab) from polylactic acid (PLA) filament. Together with a 75 mm × 50 mm cover slip (Carl Roth) these components were sonicated in ethanol for 10 min and blow dried. Afterwards, the PLA ring was glued onto the glass cover slip using two component glue (eco-sil, Picodent) and the petri dish placed inside of it. A small ridge on the edge of the bottom of the petri dish causes a ~ 200 µm high spacing between the glass slide and the bottom of the petri dish, creating a chamber.

Unstimulated PBMCs and *in vitro*-expanded, activated T cells were mixed in different ratios and resuspended in 250 µL Dulbecco’s Modified Eagle Medium (DMEM, Gibco) with 83 g L^*−*1^ of glycidyl methacrylate derivatized dextran (Dex-GMA), 5 g L^*−*1^ of LAP and 2 g L^*−*1^ of tartrazine. The suspension was pipetted into the chamber through the hole in the petri dish, and cells were allowed to settle for 10 min.

Focus support points for cells in the chamber were then set in our software. Cells were sorted using the following image analysis pipeline in our software: Detection: Brightfield; Segmentation: Contours; Filter: Size; Process: Diameter Exclusion. The size cut off in the diameter exclusion function was set to 51.06 px (equivalent to 11.5 µm). Imaging and encapsulation was conducted with the 20× objective (Plan-Apochromat 20x/0.8 M27 Carl Zeiss AG) and the 5× objective (Plan-Apochromat 5x/0,16 M27, Carl Zeiss AG), respectively, in sequential mode with 12 sections and the verify focus option. The exposure time for photopolymerization in each tile was 750 ms.

After encapsulation, the petri dish lid was removed, cells were rinsed out of the chamber, the suspension filtered through a 20 µm cell strainer (Greiner Bio-One), and isolated cells collected in the filtrate.

### Flow cytometry analysis

After sorting, cells were washed in PBS and transferred into a 96-well U-bottom plate for staining. For viability assessment, cells were stained in 50 µL BD Horizon fixable viability stain 700 (BD Biosciences) pre-diluted 1:1000 in PBS for 10 min at room temperature (RT). The cells were then washed with 150 µL FACS buffer: PBS supplemented with 0.5 wt% bovine serum albumin (BSA, Sigma-Aldrich, pre-filtered through a 0.22 µM filter (Merck)) and 2 mM ethylenediaminetetraacetic acid (EDTA, Genaxxon Bioscience). Fc-receptor blocking was performed by incubating the cells in 50 µL human Fc block reagent (BD Biosciences) pre-diluted 1:20 in FACS buffer for 10 min at RT. After washing with FACS buffer, extracellular staining was performed in 50 µL for 20 min at 4 °C with following fluorescently labelled antibodies pre-diluted in FACS buffer: anti-CD3 BV786 (1:250, BD Biosciences), anti-CD11b BB700 (1:20, BD Biosciences), anti-CD69 APC-H7 (1:10, BD Biosciences) and anti-CD137 APC (1:5, BD Biosciences). After a final washing step, cells were resuspended in FACS buffer for acquisition at the FACSLyric (BD Biosciences). Data analysis and visualization was performed using the FlowJo V.10.8.2 (FlowJo LLC) and GraphPad Prism V.10 software.

## Acknowledgements

We thank Inga Ulusoy from the Scientific Software Center of Heidelberg University for her support with software development. This work was supported by the Health + Life Science Alliance Heidelberg Mannheim, the Flagship Initiative Engineering Molecular Science of Heidelberg University within the Spotlight Project Synthetic Immunology and the Cluster of Excellence SynthImmune (to M.P. and K.G.), the Human Frontiers Science Program (RGPO03I2023, to K.G.), the Alfried Krupp Förderpreis (to K.G.), the ERC Starting Grant “ENSYNC” (No. 101076997, to K.G.) and the Carl Zeiss Foundation (to T.A.).

## Conflict of Interest

S.M., T.A. and K.G. are named inventors on a patent by the Max Planck Society that covers parts of the technology described.

## References

[1] Bonner, W. A., Hulett, H. R., Sweet, R. G. & Herzenberg, L. A. Fluorescence activated cell sorting. Review of Scientific Instruments 43, 404–409 (1972). URL 10.1063/1.1685647.

[2] LaBelle, C. A., Massaro, A., Cortés-Llanos, B., Sims, C. E. & Allbritton, N. L. Image-Based Live Cell Sorting. Trends in Biotechnology 39, 613–623 (2021). URL https://linkinghub.elsevier.com/retrieve/pii/S0167779920302705.

[3] Nitta, N. et al. Raman image-activated cell sorting. Nature Communications 11, 3452 (2020). URL https://www.nature.com/articles/s41467-020-17285-3.

[4] Nawaz, A. A. et al. Intelligent image-based deformation-assisted cell sorting with molecular specificity. Nature Methods 17, 595–599 (2020). URL https://www.nature.com/articles/s41592-020-0831-y.

[5] Isozaki, A. et al. Intelligent image-activated cell sorting 2.0. Lab on a Chip 20, 2263–2273 (2020). URL https://xlink.rsc.org/?DOI=D0LC00080A.

[6] Herbig, M. et al. Label-free imaging flow cytometry for analysis and sorting of enzymatically dissociated tissues. Scientific Reports 12, 963 (2022). URL https://www.nature.com/articles/s41598-022-05007-2.

[7] Schraivogel, D. et al. High-speed fluorescence image–enabled cell sorting. Science 375, 315–320 (2022). URL https://www.science.org/doi/10.1126/science.abj3013.

[8] Salek, M. et al. COSMOS: a platform for real-time morphology-based, label-free cell sorting using deep learning. Communications Biology 6, 971 (2023). URL https://www.nature.com/articles/s42003-023-05325-9.

[9] Chien, M.-P., Werley, C. A., Farhi, S. L. & Cohen, A. E. Photostick: a method for selective isolation of target cells from culture. Chem. Sci. 6, 1701–1705 (2015). URL 10.1039/C4SC03676J.

[10] Ungai-Salánki, R. et al. Automated single cell isolation from suspension with computer vision. Scientific Reports 6, 20375 (2016). URL https://www.nature.com/articles/srep20375.

[11] Brasko, C. et al. Intelligent image-based in situ single-cell isolation. Nature Communications 9, 226 (2018). URL https://www.nature.com/articles/s41467-017-02628-4.

[12] Nelep, C. & Eberhardt, J. Automated rare single cell picking with the als cellcelector™. Cytometry Part A 93, 1267–1270 (2018). URL https://onlinelibrary.wiley.com/doi/abs/10.1002/cyto.a.23568.

[13] Paul, R. et al. Versatile image-assisted cell sorting by selective trapping with spatiotemporal multiparameter targeting. ACS Sensors 10, 6714–6723 (2025). URL 10.1021/acssensors.5c01433.

[14] Oldenhof, S., Mytnyk, S., Arranja, A., de Puit, M. & van Esch, J. H. Imaging-assisted hydrogel formation for single cell isolation. Scientific Reports 10, 6595 (2020). URL 10.1038/s41598-020-62623-6.

[15] Hasle, N. et al. High-throughput, microscope-based sorting to dissect cellular heterogeneity. Molecular Systems Biology 16, e9442 (2020). URL https://www.embopress.org/doi/abs/10.15252/msb.20209442.

[16] Kuhn, T. M., Paulsen, M. & Cuylen-Haering, S. Accessible high-speed image-activated cell sorting. Trends in Cell Biology 34, 657–670 (2024). URL https://linkinghub.elsevier.com/retrieve/pii/S0962892424000941.

[17] Ginhoux, F., Yalin, A., Dutertre, C. A. & Amit, I. Single-cell immunology: Past, present, and future. Immunity 55, 393–404 (2022). URL https://linkinghub.elsevier.com/retrieve/pii/S1074761322000863.

[18] Patel, K. K., Tariveranmoshabad, M., Kadu, S., Shobaki, N. & June, C. From concept to cure: The evolution of CAR-T cell therapy. Molecular Therapy 33, 2123–2140 (2025). URL https://linkinghub.elsevier.com/retrieve/pii/S1525001625001790.

[19] Waugh, R. E., Lomakina, E., Amitrano, A. & Kim, M. Activation effects on the physical characteristics of t lymphocytes. Frontiers in Bioengineering and Biotechnology Volume 11 - 2023 (2023). URL https://www.frontiersin.org/journals/bioengineering-and-biotechnology/articles/10.3389/fbioe.2023.1175570.

[20] Lindner, K. et al. ESPEC-SUIT: a versatile and robust platform to identify and track antigen-specific T cell receptors in patients with cancer. Journal for ImmunoTherapy of Cancer 13, e012216 (2025). URL https://jitc.bmj.com/lookup/doi/10.1136/jitc-2025-012216.

[21] Wang, S. Y., Mak, K. L., Chen, L. Y., Chou, M. P. & Ho, C. K. Heterogeneity of human blood monocyte: two subpopulations with different sizes, phenotypes and functions. Immunology 77, 298–303 (1992).

[22] Schindelin, J. et al. Fiji: an open-source platform for biological-image analysis. Nat. Methods 9, 676–682 (2012).

